# Profiling trait and state-like behaviour in pain modulation systems: a role for effective connectivity within endogenous pain control circuits

**DOI:** 10.1101/2025.01.30.635646

**Authors:** Sonia Medina, Sophie Clarke, Sam W. Hughes

## Abstract

Endogenous pain modulation systems can be assessed through distinct psychophysical paradigms such as conditioned pain modulation (CPM), temporal summation of pain (TSP), and offset analgesia (OA). Notably, the reliability of these measures has rarely been defined within the same participants, and measures to index the consistency of each measure across sessions and in the same participants are lacking. This study examined the test-retest reliability and intra-individual consistency of CPM, TSP, and OA and explored how CPM response status across sessions relates to neural dynamics within the descending pain modulation system. In 29 healthy participants, CPM, TSP, and OA responses were assessed across two sessions. The normalised session change index (NSCI) was introduced to evaluate consistency of each measure across sessions. Spectral dynamic causal modelling (DCM) of resting-state fMRI (rs-fMRI) investigated effective connectivity within the descending pain modulation network and its association with CPM response status. OA exhibited the highest test-retest reliability and had NSCI values closest to zero, indicating stable responses. CPM and TSP showed poor reliability and NSCI values deviating from zero, reflecting greater variability. Spectral DCM analysis revealed that effective connectivity within the descending pain modulation system explained CPM response variability across sessions. Specifically, consistent strong facilitatory or inhibitory CPM was associated with greater excitatory or inhibitory PAG-to-AI effective connectivity, respectively. These findings suggest that healthy participants either demonstrate stable (i.e. trait-like) or dynamic (i.e. state-like) endogenous pain modulation across repeated sessions which can be explained based on effective connectivity within the descending pain modulation system.

## Introduction

Conditioned pain modulation (CPM), temporal summation of pain (TSP), and offset analgesia (OA) are psychophysical paradigms used to assess distinct pro- and anti-nociceptive processes [2; 17-19]. These paradigms are often used in experimental and clinical studies as biomarkers to help gain insight into dysfunctional pain mechanisms or treatment outcomes [48]. However, considerable intersession variability suggests endogenous pain modulation systems likely reflect a dynamic state of pain inhibition or facilitation in some participants [24; 48]. Being able to profile this individual between-session variability could help to identify those with dynamic (i.e. state-like) or stable (i.e. trait-like) activity across different endogenous pain modulation systems. It could also allow us to gain insight into the underlying neural dynamics that underpin the variability that is often seen across repeated psychophysical test sessions [20; 26].

Resting state functional magnetic resonance imaging (rs-fMRI) can be used to help understand variability in psychophysical measures of endogenous pain modulation [24; 27; 35]. Dynamic causal modelling (DCM) is a Bayesian framework used to infer and quantify effective connectivity, which is the directed causal influence one neural region exerts on another within a brain network [9; 10]. Unlike functional connectivity, which reflects statistical dependencies (e.g., correlations) between time-series data, DCM explicitly models how neuronal activity in one region causally influences another based on underlying biophysical principles [37]. Spectral DCM, specifically developed for rs-fMRI, leverages frequency-domain data to estimate intrinsic connectivity patterns and to understand how these relate to behavioural measures [51]. This makes DCM particularly suited for understanding how specific neural circuits are related to psychophysical measures of descending pain modulation [30; 32].

Key regions implicated in the descending pain modulation system include the periaqueductal grey (PAG), anterior cingulate cortex (ACC) and anterior insula (AI) [23; 28; 43; 44; 46]. These regions form critical nodes in top-down analgesic pathways, regulating nociceptive input through cognitive and affective influences over descending projections the spinal cord [16; 46]. By examining the directionality and strength of connectivity within this network, DCM can be used to help identify neural patterns that differentiate consistent strong inhibitors or facilitators in psychophysical paradigms which typically show high levels of intersession variability [51].

In this study, we provide a mechanistic framework to explore the neural basis of variability in endogenous pain modulation. Here, we first quantify the between-session reliability and intra-individual consistency across three distinct psychophysical endogenous pain modulation paradigms measured in the same participants (i.e. CPM, OA, TSP). We then use this information to demonstrate how DCM applied to rs-fMRI data can be used to help understand endogenous pain modulation inter-session response status, using CPM as an example.

## Methods

### Study participants

A total of 32 healthy participants were recruited to the study. Eligibility criteria included: aged 18 years or older, no history of neurological conditions or brain surgery, no psychiatric conditions, epilepsy, acute migraines, concussion which resulted in loss of consciousness, dizziness or motion sickness, no history of substance, drug or alcohol abuse, no chronic pain conditions that would impair the assessment of pain due to stimulation, not taking any medications which might affect temperature sensitivity (e.g., tricyclic antidepressants), endogenous analgesia (e.g., duloxetine) and cardiovascular medications (e.g., beta-adrenergic blocking agents), no history of hand/thumb trauma or neurological conditions affecting the hands or temperature control (e.g., Raynaud’s Syndrome) and no MRI contraindications (e.g., pregnancy, metal in body, pacemaker, etc). Participants were instructed to abstain from alcohol and recreational drugs 24 hours prior to each visit, avoid painkillers or paracetamol for 12 hours before sessions, and limit to one caffeinated drink in the morning. At the start of each session, compliance with lifestyle guidelines was checked and the experimenter confirmed that participants were not in any pain at the time of the session. Any deviations led to immediate termination and rescheduling. All participants provided written, informed consent at the beginning of their first visit. The study was approved by the Health Research Authority and Health and Care Research Wales ethics committee (Ethics reference: 22/HRA/4672). Participants were informed that they could withdraw from the study at any time during informed consent process.

### Study design

Participants attended five sessions in total as part of a wider study (for full details, refer to the study preregistration (doi.org/10.17605/OSF.IO/X9UF8) and [31]. In the current report, procedures carried out as part of sessions 1-4 are presented. In order to assess nociceptive and pain modulatory function, three protocols were employed: conditioned pain modulation (CPM), temporal summation of pain (TSP) and offset analgesia (OA). All three protocols were repeated across two sessions in order to examine their test-retest reliability. The CPM and TSP protocols were conducted sequentially, with CPM immediately following TSP, at the beginning of two out of the three first visits (i.e., either on visits 1 and 2, 1 and 3 or 2 and 3). The OA protocol was completed at the end of two out of the three visits. CPM/TSP and OA measurements were not acquired within the same visits for all participants (for example, some participants completed CPM/TSP at the beginning of visits 1 and 2, and OA at the end of visits 2 and 3). This variability depended on availability of laboratory and equipment resources. The experimenter was kept consistent across all sessions. All test-retest sessions were conducted at the same time of the day, approximately one week apart (average days between sessions = 9.4 SD = 6). The MRI visit (session 4) was scheduled between 2-30 days apart from the other visits (Figure 1).

**Figure 1.**
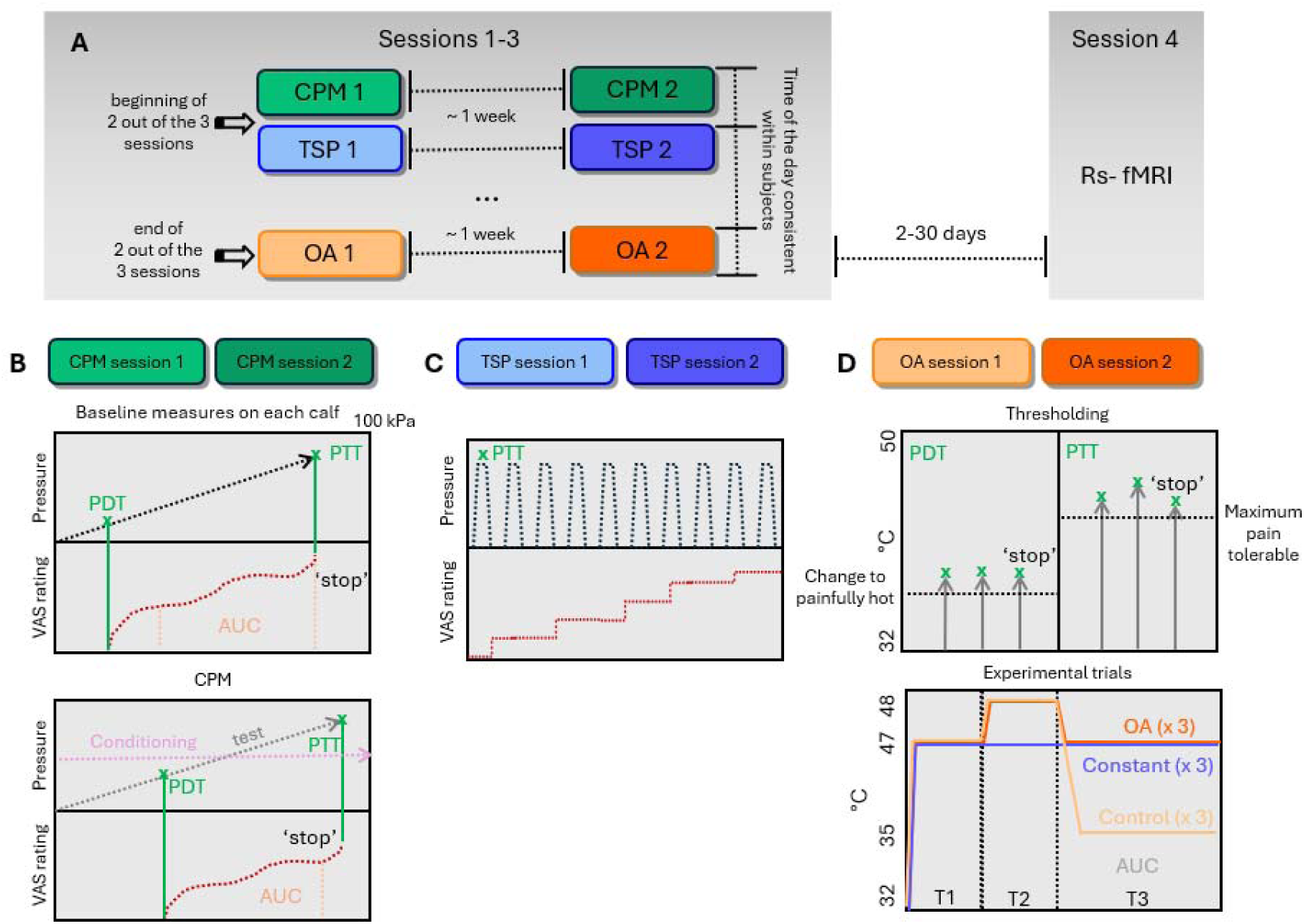
Study design and procedures. Participants attended 3 sessions approximately one week apart and at the same time of the day. At the beginning of two of these three sessions, participants completed identical CPM and TSP protocols (N=29). For CPM (lower left panel) participants rated a steadily increasing pressure stimulus in each of the calves with a VAS. The pressure at which they first reported pain was their pain detection threshold (PDT) and the pressure at which participants pressed the stop button was their pain tolerability threshold (PTT). For CPM, participants rated an identical pressure ramp on their right calf while they experience tonic pressure on their left calf, individually calibrated at 70% their PTT on their left calf. CPM was defined as the difference in PDT and PTT from baseline to CPM runs, as well as the difference in the area under the curve (AUC) between runs within the painful pressure range overlapping across baseline and CPM runs. For TSP (lower middle panel), ten consecutive pressure stimuli were delivered to participants’ right calves, and they were instructed to rate them on a VAS. At the end of two out of three sessions (not necessarily overlapping fully with CPM and TSP measurements) participants completed identical OA protocols (N=25, lower right panel) following procedures described by Grill and Coghill (2002). For heat pain detection thresholding, three heat ramps were delivered to participants’ left forearm, and participants were instructed to press the button as soon as they felt pain. PDT resulted from averaging the temperatures at which participants pressed the button. For PTT, three heat ramps were delivered to an adjacent location of the left forearm, and participants were instructed to press the button as soon as they felt ‘moderate pain’. PTT resulted from the average temperature at which participants pressed the button. Participants complete an fMRI session in a different visit.

### Study Procedures

#### Conditioned Pain Modulation (CPM) and temporal summation of Pain (TSP)

CPM and TSP protocols were conducted with computer-controlled cuff-algometry system (LabBench software 4.7.3 and CPAR+ device, Aalborg, Denmark). Participants sat on a padded armchair with leg support raised up to approximately 45 degrees. Independent computerised torniquet cuffs were placed around each calf (10 x 61 cm) [6; 13]. Test stimulus and conditioning stimulus were always applied to the right and left calves, respectively. Participants were handed a computerised visual analogue scale (VAS) anchored from ‘no pain’ to ‘maximal pain’ and a slider set to ‘no pain’ as a default.

First, the test stimulus cuff began to inflate at a rate of 1 kPa/sec to a maximum of 100 kPa. Participants were instructed to move the slider on the computerised VAS once they felt their first pain sensation, and continuously rate their pain until the maximum they could tolerate. Once they reached their maximum tolerable pain, participants could push a ‘stop’ button placed next to the computerised VAS, which deflated the cuff immediately. The pressure at which participants first moved the VAS slider was considered their pain detection threshold (PDT), and the pressure registered immediately before participants pressed the ‘stop’ button was considered their pain tolerability threshold (PTT). If participants did not press the ‘stop’ button before reaching the maximum pressure, then this was considered their PTT (i.e., 100 kPa). Following this, an identical examination was performed on the conditioning stimulus cuff.

Following a 2-minute resting period, the TSP protocol was carried out on the test stimulus calf; ten 1-second stimuli with pressure set to participants individual PTT recorded during the earlier assessment on the same calf. The cuff inflated and deflated rapidly for each stimulation. Stimuli had a 2-second duration with 1-second interval between them. Participants were instructed to rate their perceived pain intensity on each stimulus using the computerised VAS without returning it to ‘no pain’ between stimuli.

Following a 5-minute resting period, the final part of the CPM protocol was carried out. The conditioning stimulus cuff (left calf) inflated rapidly to a pressure equal to 70% participants’ PTT on the same calf, and it remained inflated until the end of the protocol to serve as concurrent conditioning tonic stimulus. Once it reached the target pressure, the test stimulus cuff started to inflate at a rate of 1 kPa/sec. Participants were instructed to rate the test stimulus (right calf) in the exact same way as they did at the beginning. Participants’ PDT, PTT and continuous VAS ratings were recorded again.

#### Offset Analgesia (OA)

Heat stimuli were delivered to the volar surface of the dominant forearm via a 30x 30mm Peltier device (Medoc TSAII, Ramat Yishai, Israel) attached with a Velcro strap. The baseline temperature was set at 32°C. For OA trials, temperature-rise/fall rates were set at 6°C/s. When constant rating was required, participants rated their pain using a computerised visual analogue scale (VAS) with verbal anchors of “no pain sensation” and “worst pain imaginable” displayed on a computer screen located in front of participants. The cursor was moved across the scale by participants using the keyboard. OA testing was divided in three phases:

1. *Familiarisation phase.* One single series of eight temperature ramps was delivered on the volar surface of the dominant forearm. Each of these ramps consisted of a gradually increasing stimulus from baseline (32°C) to one of the target temperatures, staying constant for 5 seconds and decreasing back to baseline. Target temperatures were, 35°C, 43°C, 44°C, 45°C, 46°C, 47°C, 48°C, and 49°C, in random order. Participants were instructed to constantly rate their perceived pain for each stimulus through a VAS.
2. *Thresholding phase*. Three consecutive ascending temperature ramps from baseline at a rate of 1°C/second were delivered to the same forearm location. Participants were instructed to click on a mouse as soon as they felt the first painful sensation. The resulting average temperature at which participants clicked the mouse was considered their heat PDT. For heat PTT’s, identical temperature ramps were delivered, and participants were instructed to click on the mouse as soon as they felt ‘moderate pain’, specified as an 80/100 on a numerical rating scale from 0 (‘no pain’) to 100 (‘worst pain’). The resulting average temperature at which participants clicked the mouse was considered their heat PTT. For safety purposes, maximum temperature of 52°C was reachable. In those cases where participants reached the maximum, 52°C was considered the PTT temperature.
3. *Experimental phase.* The experimental trials followed the paradigm proposed by Grill and Coghill (2002), comprising three conditions and three trials per condition (Figure 2): the *OA condition* consisted of a train of three consecutive temperatures (47°C-48°C-47°C) with a duration of 5 seconds for temperature one (T1), 5 seconds for temperature two (T2) and 20 seconds for temperature three (T3). OA effect was expected to be triggered after the 1-degree decrease from T2 to T3; the *control condition* was similar to the OA condition, however, the temperature during T3 was 35°C (i.e. 47°C-48°C-35°C); the *constant condition* consisted of a constant 47°C heat stimulus with a duration of 30 seconds. During stimulation, participants continuously rated the intensity of their perceived pain through a computerised VAS. There was a one-minute pause between each trial, and the trial order was pseudo-randomised across participants. In order to avoid any kind of sensitisation effect, each stimulus was administered on a different but adjacent part of the volar surface of participants’ dominant forearms.

**Figure 2.**
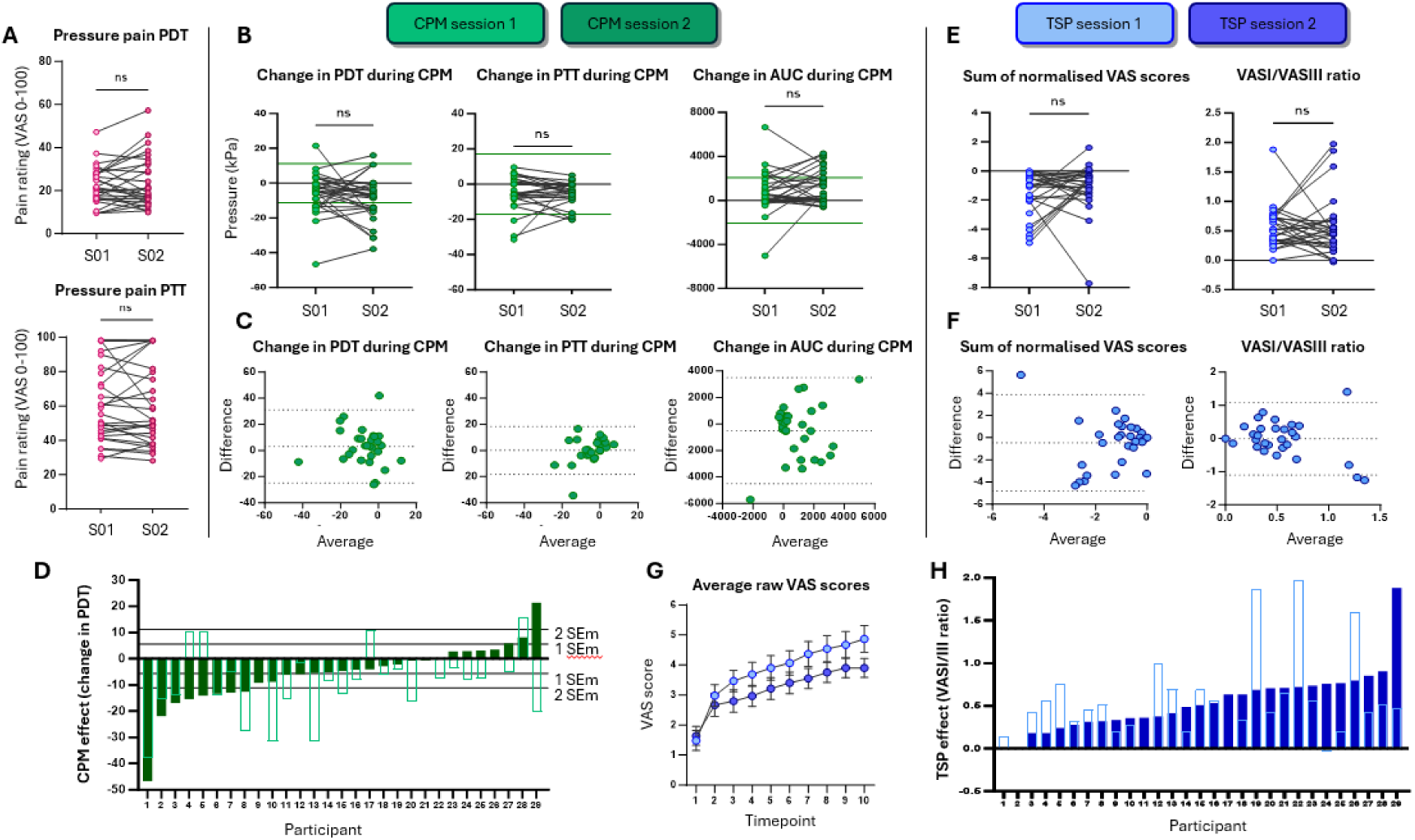
Summary of results from CPM and TSP protocols. Pressure PDT and PTT results are depicted on the left (pink). A (top panel): delta values for PDT, PTT and AUC within each session, representing the CPM effect (top left three panels, green) and TSP results within each session (top right three panels, blue). B (middle panel): Bland-Altman plots representing the relationship between average effects across session and the difference between session for each individual. For change in PDT during CPM, bias=3.156 (SD=14.25), 95% limits of agreement (LoA), −24.78 to 31.09. For change in PTT during CPM, bias=0.5793 (SD=10.61), 95% LoA, −20.22 to 21.38. For change in AUC during CPM, bias=-502.4 (SD=2042), 95% LoA, −4505 to 3500. For sum of normalised VAS scores during TSP, bias=-0.4692 (SD=2.215), 95% LoA, −4.81 to 3.872. For VASI/VASIII ratio, bias=-0.0026 (SD=0.5588), 95% LoA, −0.1098 to 0.1092. C (bottom panel): CPM effect (absolute change in PDT, lower left panel) plotted for each individual participant in order of magnitude, from largest inhibitory response (left) to largest facilitatory response (right), for session 1 (dark green solid bars). The responses from session 2 for the same participants are then overlaid (pale green open bars), showing the variability in some individuals between the two sessions. Responders to CPM were defined as those with difference outside 2 standard error of measurement (SEM). TSP effect (VASI/III ratio, lower right panel) also plotted for each individual participant in order of magnitude, from smallest response (left) to largest response (right), for session 1 (dark blue solid bars). The responses from session 2 for the same participants are then overlaid (pale blue open bars), showing the variability in some individuals between the two sessions.

**Figure 3.**
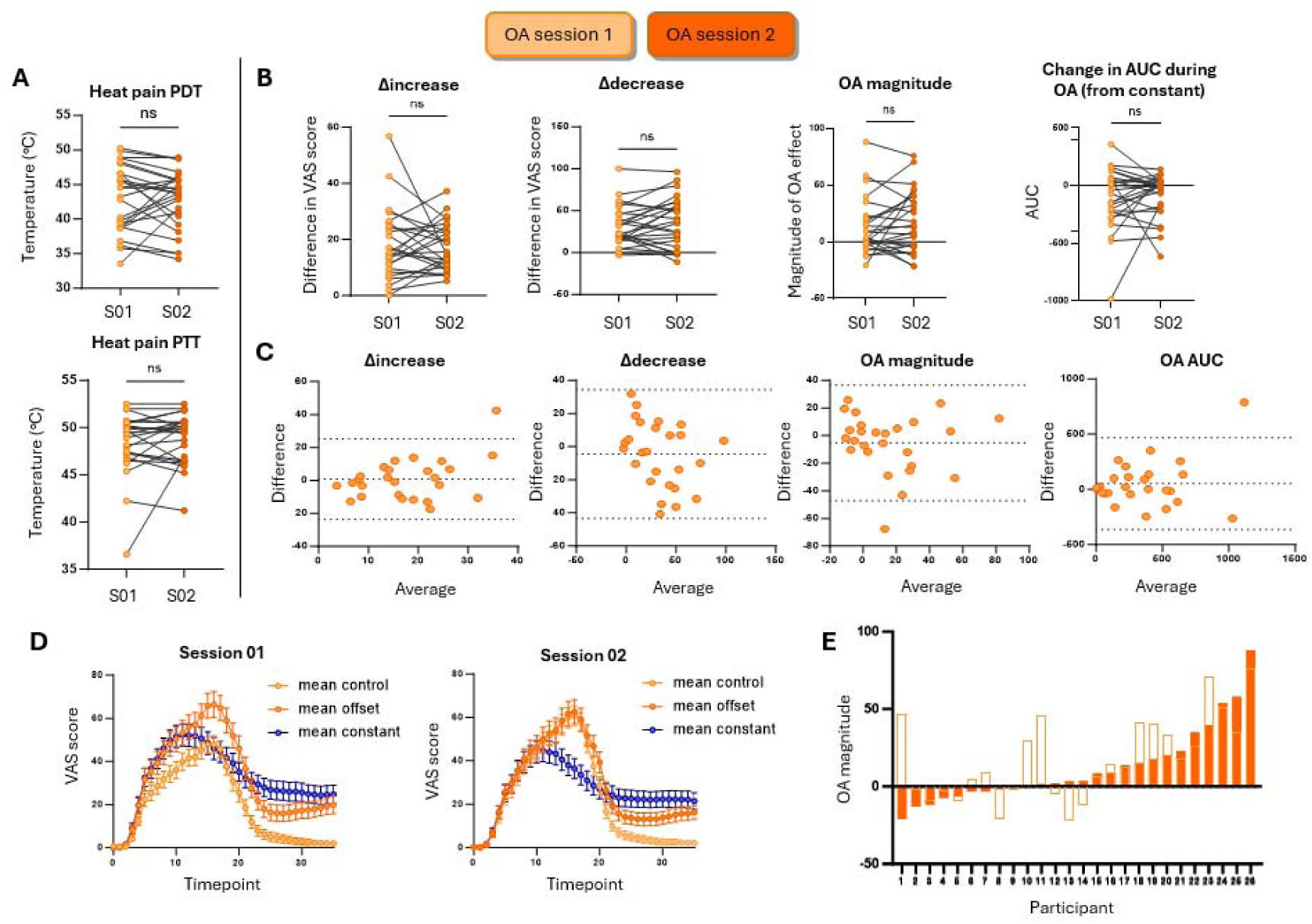
Summary of results for OA protocol. Heat PDT and PTT are shown on the left panel. A (top panel): effect of increase and decrease of temperature for OA trials (averaged across three runs), OA magnitude and OA AUC measures were not significantly different between session 01 and 02 (paired t-test p=0.7402, p=0.2617, p=0.2179 and p=0.2164, respectively). B: Bland-Altman plots. For OA effect of increase in temperature, bias=0.8203 (SD=12.48), 95% LoA, −23.64 to 25.28. For OA effect of decrease in temperature, bias=-4.466(SD=19.83), 95% LoA, −43.33 to 34.40. For OA magnitude, bias=-5.286 (SD=21.32), 95% LoA, −47.08 to 36.51. Lastly, for OA AUC, bias=52.76 (SD=212.1), 95% LoA, −363 to 468.6. C: Mean responses from all participants plotted for session 1 and 2. Error bars represent the SEM.D: OA magnitude plotted for each individual participant in order of magnitude for session 1 (dark orange solid bars). The responses from session 2 for the same participants are then overlaid (pale orange open bars), showing the variability in some individuals between the two sessions.

#### Functional Magnetic Resonance Imaging (fMRI)

Participants completed a ∼6-minute resting state fMRI scan during immersion of their right hand in gelled-cold water as a close in-scanner alternative to the cold pressor task (CPT). This strategy successfully induced variable levels of tonic mild-to-moderate pain across participants. For a full report of design, materials and data preprocessing see Medina and Hughes (2024) [32].

### Statistical analysis

All data analyses described below, except for those specified as *exploratory* in the section title, are part of preregistered analysis plan (doi.org/10.17605/OSF.IO/X9UF8) CPM, TSP and OA measures were obtained via a custom-made script in MathWorks^R^ Matlab R2023b. All reliability analyses were performed on IBM SPSS Statistics 29.0. Statistical significance was set at *p* < 0.05.

#### CPM effect

Within each session, two strategies to calculate CPM effects were followed. On one hand, CPM was assessed as a change in pain *thresholds* during tonic stimulation. This is, CPM measures, calculated separately for PDT and PTT, resulted from the absolute difference in pain thresholds of the test stimulus during baseline and conditioning tests (i.e., CPM = pain threshold _baseline_ – pain threshold _conditioning_). Here, positive and negative CPM values reflected descending facilitation and inhibition, respectively. CPM was also assessed as a change in overall pain *ratings* throughout the trials. CPM measures resulted from the absolute difference in the area under the curve (AUC) for continuous VAS ratings of the test stimulus within the overlapping range across baseline and conditioning tests for each run (i.e., CPM = AUC _baseline_ – AUC _conditioning_; Figure 1). Here, positive and negative CPM values reflected descending inhibition and facilitation, respectively. Finally, paired-sample t-test were performed for each CPM measure of interest in order to examine differences across sessions.

#### Participant stratification

For each CPM measure of interest (i.e., delta PDT, delta PTT and delta AUC), the standard error of measurement (SEM) was calculated as a measure of absolute reliability or consistency and to distinguish change greater than measurement error [6; 21]. SEMs were calculated as follows: SEM = standard deviation of baseline measure × √ (1−(ICC of baseline measure)), where ‘baseline measure’ refers to the relevant baseline value for each parameter of interest and ICC corresponds to the interclass correlation coefficient (ICC (2, k)). For each measure of interest, participants showing a CPM response outside of the ± 2SEM range are considered CPM *responders*, which could show either a facilitatory (positive values) or an inhibitory CPM response. Participants showing a CPM response within the ± 2SEM range are referred to as CPM *non-responders*.

#### TSP effect

Within each session, mean VAS scores during the pauses between each of the 10 stimulations were extracted for analysis. Each score was normalised by subtracting the mean VAS score following the first stimulation. TSP was assessed via two strategies; firstly, TSP measures resulted from the sum of all the normalised VAS scores, indicating whether participants generally rated subsequent stimuli as more or less painful compared to the first. Here, negative values indicated the presence of TSP. Secondly, the mean values of the first three normalised VAS scores (VAS-I) and the last three normalised VAS scores (VAS-III) were calculated, and TSP measures resulted from the VAS-I/VAS-III ratio. Here, positive values indicated the presence of TSP. Again, paired-sample t-test were performed for each TSP measure of interest in order to examine differences across sessions.

#### OA effect

For calculation of OA effect within each session, a number of measures were computed; within each condition, to calculate the effect of an increase of temperature in perceived pain intensity (Δ*increase*), the absolute difference between the VAS rating reported four seconds after the beginning of T1, and the VAS rating reported four seconds after the beginning of T2 was calculated. Similarly, the absolute difference in VAS rating between second 4 of T2 and second 4 of T3 represented the change in perceived pain intensity after a decrease of temperature (Δ*decrease*). These calculations were also carried out for the constant condition as control measures. The difference between Δdecrease and Δincrease in the OA condition is considered a relative measure of the magnitude of the OA effect (*OA magnitude*). Here, higher and lower values indicated higher and lower OA magnitude, respectively. Possible main effects of Condition and Session, as well as interaction effects between these two, were assessed separately for Δincreases and Δdecreases using a two-way repeated measures analysis of variance (ANOVA), with session and condition as within-subjects factors. Δincreases and Δdecreases were also compared within each condition via paired samples t-tests. The effect of Session on the OA magnitude was examined via a paired-samples t-test.

In addition, the AUC for VAS scores throughout T3 were calculated for each trial and averaged across the three trials per condition. OA was also assessed as the difference between AUC during the OA condition and AUC during the Constant condition (i.e., OA_AUC_ = AUC_T3 OA_ – AUC_T3 Constant_). Here, negative and positive values indicate presence or absence of OA_AUC_ effect, respectively. Differences in CPM_AUC_ across sessions were examined via a paired-samples t-test. All post-hoc pairwise comparisons were corrected for multiple comparisons using a Holm Sidak’s adjustment.

#### Test-retest reliability

Reliability of all measures of interest across CPM and TSP and OA protocols were assessed with the intraclass correlation coefficient (ICC (2, k)) [45]. Reliability was interpreted according to the following ICC ranges: poor < 0.4; fair 0.4–0.59; good 0.6–0.74; excellent >0.75 [39].

#### Exploratory analysis: Normalised Session Change Index (NSCI)

We proposed a novel way of assessing the consistency of pain modulation measures (i.e., CPM, TSP and OA) across sessions, with the aim of providing a standardised measure of change and facilitate comparisons across measures or datasets. The Normalised Session Change Index (NSCI) quantifies the relative change between two sessions normalised by the standard deviation of the first session. NSCIs were calculated for each one of our measures of interest (CPM_PDT_, CPM_PTT_, CPM_AUC_, TSP, TSP_VASI/VASIII_, OA_Δincrease_, OA_Δdecrease_, OA_magnitude_, OA_AUC_) via computing the difference between session 1 and 2, divided by the standard deviation of session 1 across the whole sample. Therefore, lower NSCI absolute values indicated greater consistency of measures across sessions, and higher NSCI absolute values indicated greater relative volatility, suggesting significant changes between the sessions relative to the variability observed in the first session. Positive NSCI values indicated that the measure of interest was higher in session 1 than in session 2. Negative NSCI values indicated that the measure of interest was higher in session 2 than in session 1.

#### Exploratory analysis: fMRI

We set out to explore whether variability in CPM responses across sessions could be predicted by specific patterns of effective connectivity among areas that previously showed a significant engagement during tonic pain in the same individuals, namely the dorsal anterior cingulate cortex (dACC) bilaterally, the anterior insula (AI) bilaterally, the thalamus bilaterally and the periaqueductal grey (PAG). A full report of the results, as well of region of interest description can be found in Medina and Hughes (2024) [32]. To do this, we performed spectral dynamic causal modelling (spDCM) to explore the most probable patterns of excitatory and inhibitory influences from one region to another (i.e., *effective connectivity*) that underpinned variability in CPM response status across sessions.

#### CPM response status

While participants’ NSCI allowed us to quantify the standard difference across sessions (i.e., measure volability) by taking into account to the total variability in the sample in session 1, it does not provide specific information about whether participants were considered ‘responders’ or ‘non responders’ according to our pre stablished SEM criteria. In order to address this a new variable was created, where participants were assigned a single composite score referred to as their “CPM status.” This score was determined by summing their CPM response following stratification within each session, based on SEM thresholds. The scoring system was as follows: −2 for inhibitory CPM responses below −2 SEM, −1 for inhibitory CPM responses below −1 SEM, 0 for CPM responses within ±1 SEM, 1 for facilitatory CPM responses above 1 SEM and 2 for facilitatory CPM response above 2 SEM. The final CPM status ranged from −4 (indicative of strong inhibitory responses in both sessions) to 4 (indicative of strong facilitatory responses in both sessions).

For spDCM, the DCM toolbox implemented in SPM12[1] was used. First, a fully connective model was defined, with all possible connections among ROIs switched on. Model parameters were estimated through variational Laplace inversion, utilising the default prior probability densities provided by the toolbox. Effective connectivity strengths were assessed at the group level within a Parametric Empirical Bayes (PEB) framework[51]. The PEB design matrix included three regressors: the group mean, the CPM response status, and NSCI scores, which allowed to explore the effect of CPM response status accounting for the individual volatility in CPM responses across sessions (Figure 5). Only connections with a posterior probability (pp) greater than 0.99 of being present versus being absent are reported as significant. Model validation was performed using leave-one-out cross validation (LOOCV) implemented in SPM (spm_dcm_loo.m). For LOOCV, we focused on the connection with the highest absolute effective connectivity parameter, since the inclusion of many connections can increase the model complexity and therefore compromised its predictive capacity.

**Figure 4.**
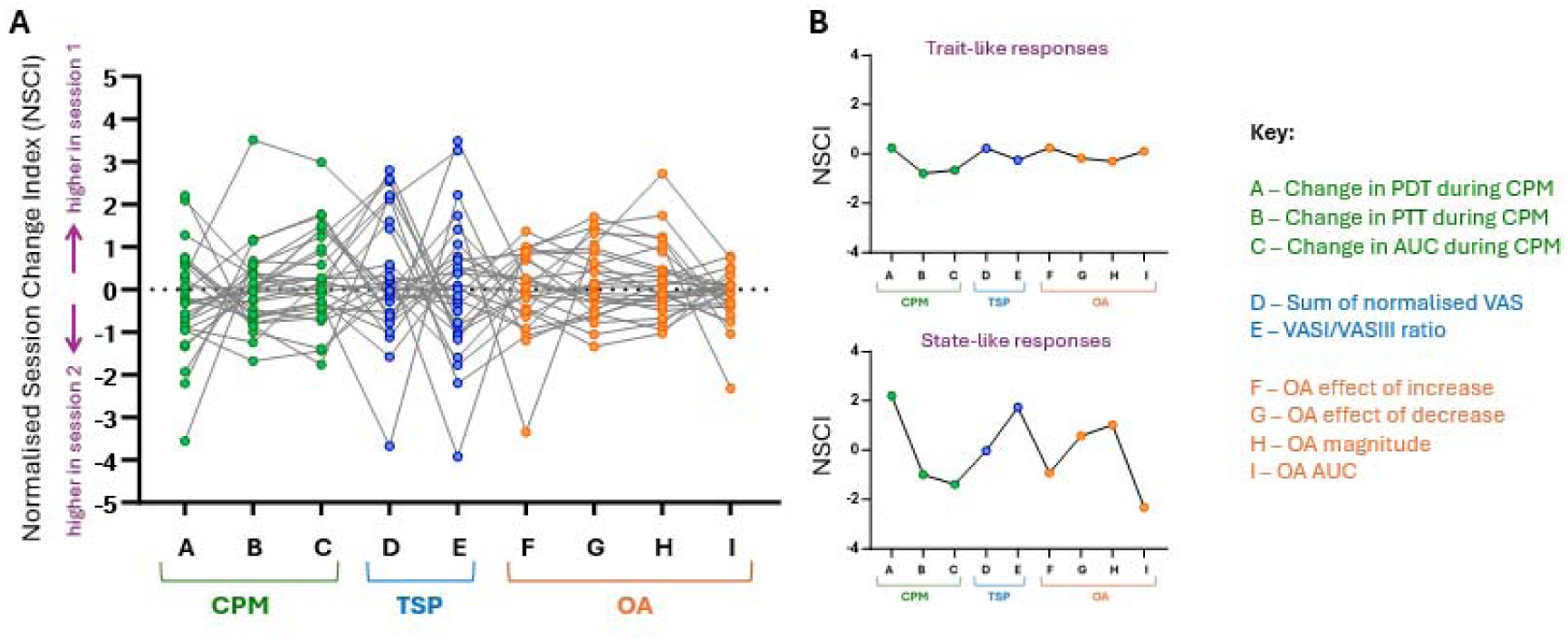
NSCI index for consistency of responses across all paradigms. A) Normalised Session Change Index (NSCI), which quantifies the relative change between the two sessions normalized by the standard deviation of the first session, is plotted for each measure. The NSCI is calculated as the difference between session 1 and 2, divided by the standard deviation of session 1. Higher NSCI values indicate greater relative volatility, suggesting significant changes between the sessions relative to the variability observed in the first session. B) NSCI data is shown for two example participants, with the first (top panel) showing very consistent responses across the two sessions and all measures, and the second (bottom panel) showing volatile responses across the measures and sessions.

**Figure 5.**
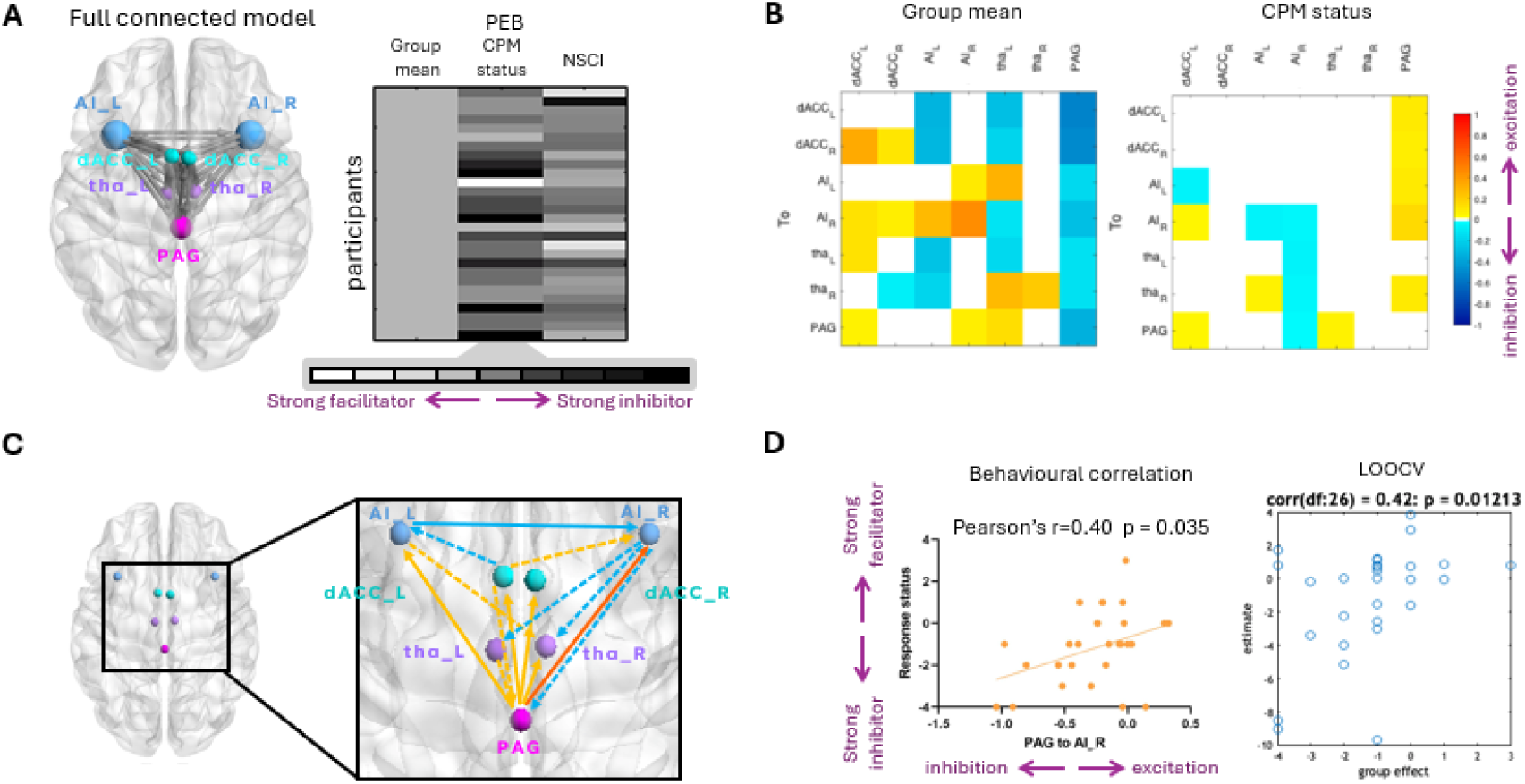
DCM results. A full-connected model was described including the dACC, AI and thalamus bilaterally and PAG. Following model parameter estimation, a PEB matrix was constructed for group analysis with group mean, CPM status and NSCI as regressors (top left panel). Light and darker shades of grey indicate high and low values, respectively. Cross correlation matrices depict effective connectivity parameters with pp>0.99 (top right panel). Effective connectivity pattern for main effect of CPM status is depicted in the lower right panel, with solid arrows corresponding with excitatory (yellow) and inhibitory (blue) bottom-up or within region connectivity and dashed arrows for excitatory (yellow) and inhibitory (blue) top-down connectivity. Orange arrow corresponds to connection selected for LOOCV (lower left panel).

## Results

Two participants withdrew from the study. One participant was excluded due to pain responses consistently reported outside the normal range (i.e, no pain reported across most tests). Therefore, 29 participants are included for analysis; mean age 29 (range: 19-58; SD = 10, 18 were females). Three participants from the final sample did not complete OA testing due to equipment availability issues at the start of the study, resulting in a sample size of 26 participants for OA.

### Baseline measures: pressure and heat pain thresholds

At group level for pressure pain thresholds, we observed no significant differences across sessions for PDTs [Session 1: mean (SD) = 22.20(8.79); Session 2: mean (SD) = 23.78(11.90)] or PTTs [Session 1: mean (SD) = 62.72(23.24); Session 2: mean (SD) = 62.02(24.62)]. We also observed not significant group differences across sessions for heat PDT [Session 1: mean (SD) = 42.92(4.90); Session 2: mean (SD) = 42.84(4.26)] and heat PTT [Session 1: mean (SD) = 48.38(3.33); Session 2: mean (SD) = 48.87(2.61)].

### CPM

For each measure of interest (CPM_PDT_, CPM_PTT_, CPM_AUC_) responses were stratified according to the SEM as follows: facilitatory response (>+2SEM), inhibitory response (<-2SEM) and non-response (within ± 2SEM; Figure 2). For CPM_PDT_, the SEM = 5.60 (2SEM = 11.20), yielding 8 inhibitors, 1 facilitator and 20 non-responders for Session 1, and 11 inhibitors, 1 facilitator and 17 non-responders for Session 2. For CPM_PTT_, the SEM = 8.55 (2SEM = 17.11), corresponding to 3 inhibitors, no facilitators and 26 non-responders for Session 1 and for Session 2. Finally, for CPM_AUC_, SEM = 1034 (2SEM = 2068, in arbitrary area units), yielding 6 inhibitors, 1 facilitator and 22 non-responders for Session 1, and 8 inhibitors, no facilitators and 21 non-responders in Session2. At group level, paired samples t-tests revealed no significant differences across sessions in the change in PDT during CPM, change in PTT during CPM or change in AUC of VAS ratings during CPM (p = 0.243, p = 0.947 and p = 0.195, respectively).

### TSP

Average raw values following each pressure stimulus reflected a progressive increase in pain with time across both sessions (Figure 2A). The overall sum of normalised VAS scores (i.e., TSP) in Session 1 indicated that 24 participants reported an overall increase in pain relative to the first stimulation (TSP < 0) and 5 participants did not report TSP (= 0). In Session 2, 25 participants reported TSP (< 0), 2 participants did not report TSP (= 0), and 2 participants reported an overall decrease in pain relative to the first stimulation (TSP > 0). TSP _ASI/VASIII_ results indicated that 27 participants reported more pain within the last three trials than on the first three trials (> 0), 1 participant reported no differences (= 0), and 1 participant reported a mild decrease in pain (< 0). In Session 2, 25 participants reported more pain within the last three trials than on the first three trials, 3 participants reported no differences (= 0), and 1 participant reported a mild decrease in pain (< 0). At group level, paired t-test revealed no significant differences across sessions for TSP or TSP_VASI/VASIII_ (p = 0.263 and p = 0.979, respectively).

### OA

For Δ*increase,* we observed a main effect of Condition (F _(2,50)_ = 110.52, p < 0.001). Post-hoc pairwise comparisons revealed that for the Constant trials was significantly lower than for OA and Control trials (all p’s<0.001). There were no significant differences between OA and Control trials (p = 0.307). There was no significant main effect of Session or interaction effects.

For Δ*decrease,* repeated measures ANOVA revealed a main effect of Condition (F _(2,50)_ = 20.886, p < 0.001) and a Session x Condition interaction effect (F _(2,50)_ = 20.886, p = 0.014). Post-hoc pairwise comparisons indicated that this interaction was like driven by Δdecrease being significantly higher in the Control condition for Session 2 than for Session 1 (p = 0.018) likely due to the fact that participants rated on average T1 and T2 on the Control condition during Session 1 considerably less painful than on Session 2. Comparisons across conditions within sessions revealed that Δdecrease was significantly lower in the Constant condition than in OA and Control conditions in both sessions (Session 1: p = 0.020 and p = 0.011, respectively; Session 2: all p<0.001) and there were no significant differences between OA and Control conditions across sessions (p = 0.999 and p = 0.134 for Session 1 and 2, respectively). There was no significant main effect of Session.

When comparing directly Δincrease Δdecrease within each condition and session, we observed a significant difference across all t-tests, where all Δincrease were lower than Δdecrease (Session 1: p=0.015, p<0.001and p<0.001 for OA, Control and Constant conditions, respectively; Session 2: p=0.002, p<0.001and p<0.001 for OA, Control and Constant conditions, respectively). Paired-samples t-test revealed no significant differences across sessions for OA magnitude and OA_AUC_ (p = 0.218 and p = 0.621 respectively).

### Test-retest reliability for CPM, TSP and OA

ICC (2, k) scores indicated that baseline pressure pain measures showed good-to-excellent reliability (all >0.70), with pressure PTT reaching the highest reliability (0.87). Pressure pain modulation measures showed poor reliability (all <0.40) with TSP ICCs particularly low (<0.2). ICC observed for heat pain thresholds indicated excellent reliability (>0.75), especially for PDT (0.79). OA measures showed also good-to-excellent reliability, where lowest ICC corresponded to OA magnitude (0.66) and highest ICC corresponded to OA AUC (0.77). A summary of ICCs is shown in Table 1.

**Table 1.** Summary of reliability results. The intraclass correlation coefficient 2, k (ICC; two-way random effects model) was calculated and interpreted using the following ranges: poor<0.4; fair=0.4–0.59; good=0.6–0.74; excellent>0.75. CI = confidence interval.

| Measure | Specific measure | ICC (2,k) | Interpretation |
| --- | --- | --- | --- |
| Baseline | Pressure pain detection threshold (PDT) | 0.71 (95% CI [0.48, 0.85]) | Good |
|  | Pressure pain tolerance threshold (PTT) | 0.87 (95% CI [0.75, 0.94]) | Excellent |
|  | Baseline VAS ratings area under the curve (AUC) | 0.75 (95% CI [0.52, 0.87]) | Excellent |
| Conditioned pain modulation (CPM) | Change in PDT during CPM | 0.34 (95% CI [0.02, 0.62]) | Poor |
|  | Change in PTT during CPM | 0.38 (95% CI [0.02, 0.65]) | Poor |
|  | Change in VAS ratings AUC during CPM | 0.30 (95% CI [-0.06, 0.60]) | Poor |
| Temporal summation of pain (TSP) | Sum of normalised VAS scores for TSP | 0.01 (95% CI [-0.35, 0.36]) | Poor |
|  | VAS-I/VAS-III ratio for TSP | 0.19 (95% CI [-0.18, 0.51]) | Poor |
| Offset analgesia (OA) | Heat pain PDT for OA | 0.79 (95% CI [0.54 – 0.90]) | Excellent |
|  | Heat pain PTT for OA | 0.77 (95% CI [0.49 – 0.90]) | Excellent |
| | OA Δdecrease | 0.73 (95% CI [0.49 – 0.87]) | Good |
|  | OA magnitude | 0.66 (95% CI [0.38 – 0.83]) | Good |
|  | OA AUC | 0.77 (95% CI [0.55–0.89]) | Excellent |

| Measure | Specific measure | ICC (2,k) | Interpretation |
| --- | --- | --- | --- |
| Baseline | Pressure pain detection threshold (PDT) | 0.71 (95% CI [0.48, 0.85]) | Good |
|  | Pressure pain tolerance threshold (PTT) | 0.87 (95% CI [0.75, 0.94]) | Excellent |
|  | Baseline VAS ratings area under the curve (AUC) | 0.75 (95% CI [0.52, 0.87]) | Excellent |
| Conditioned pain modulation (CPM) | Change in PDT during CPM | 0.34 (95% CI [0.02, 0.62]) | Poor |
|  | Change in PTT during CPM | 0.38 (95% CI [0.02, 0.65]) | Poor |
|  | Change in VAS ratings AUC during CPM | 0.30 (95% CI [-0.06, 0.60]) | Poor |
| Temporal summation of pain (TSP) | Sum of normalised VAS scores for TSP | 0.01 (95% CI [-0.35, 0.36]) | Poor |
|  | VAS-I/VAS-III ratio for TSP | 0.19 (95% CI [-0.18, 0.51]) | Poor |
| Offset analgesia (OA) | Heat pain PDT for OA | 0.79 (95% CI [0.54 – 0.90]) | Excellent |
|  | Heat pain PTT for OA | 0.77 (95% CI [0.49 – 0.90]) | Excellent |
|  | OA Δdecrease | 0.73 (95% CI [0.49 – 0.87]) | Good |
|  | OA magnitude | 0.66 (95% CI [0.38 – 0.83]) | Good |
|  | OA AUC | 0.77 (95% CI [0.55-0.89]) | Excellent |

### Intra-individual consistency for CPM, TSP and OA measured using NSCI

Overall, heat pain measures (heat PDT, PTT and OA measures) were the most consistent, with NSCIs values closest to zero (Figure 5). OA AUC measures were the most consistent, and TSP values were the most volatile, with higher range of NSCI values around zero. In general, volatility between sessions did not present on a particular direction, with some measures being higher in Session 1 (positive NSCI) and some measures being higher in Session 2 (negative NSCI) and NSCI switching directions across different measures within individuals.

### DCM and CPM response status

Individual spDCM models explained an average of 90.5% of data variability, suggesting a good data fit. PEB results for the effect of CPM status revealed that variability in this composite score was characterised by mostly bottom-up excitatory influences from the PAG to left AI, dACC bilaterally and right thalamus, and from the left dACC to the right AI. We also observed top-down excitatory influences from the left AI to the contralateral thalamus and from the left thalamus to the PAG. Regarding inhibitory influences, the most probable inhibitory connections were observed top-down from the right AI to thalamus bilaterally and PAG, and also from the left dACC to the ipsilateral AI and from left to right AI. For LOOCV, we included the connection with the highest absolute connectivity weight, namely the connection from the PAG to the right AI. The predicted CPM status correlated significantly with the actual CPM status (r = 0.42, p = 0.012). To facilitate the interpretability of these results, we performed Pearson’s correlation between the individual PAG to right AI effective connectivity parameters and CPM status, which indicated that there was a significant positive correlation (r = 0.40, p = 0.035). This is, participants with high excitatory effective connectivity from the PAG to the right AI were strong CPM facilitators across both sessions, and vice versa.

## Discussion

This study examined the between-session reliability of CPM, TSP, and OA in healthy participants. We also introduce a novel approach for classifying participants according to the magnitude and consistency of their CPM response across sessions using SEM thresholds (e.g., strong inhibitors, facilitators, or neutral responders) and NSCI to distinguish between stable (close to zero NSCI) and variable (high or low NSCI) measures, respectively. We then integrated this framework into spectral DCM to investigate how neural connectivity patterns within the descending pain modulation network relate to these psychophysical CPM profiles. Using this approach, we show a correlation between PAG-to-AI effective connectivity and CPM response status which suggests that intrinsic neural dynamics in endogenous pain modulatory pathways can predict consistent inhibitory or facilitatory profiles. These results highlight the potential of using effective connectivity parameters as biomarkers for understanding individual stable (trait-like) versus dynamic (state-like) endogenous pain modulation profiles.

Our findings demonstrate notable differences in the reliability of CPM, TSP, and OA measures. Consistent with previous studies, OA exhibited good-to-excellent reliability across sessions, which likely reflects a more robust localised brainstem mechanism reflecting the temporal filtering of nociceptive signals [17; 49]. In contrast, CPM and TSP showed poor reliability, highlighting the inherent variability often seen in these paradigms [12]. These differences may stem from there being cortical influences over descending pain modulation pathways assessed by CPM [50]. Variability in CPM measures was particularly evident when stratifying participants into responders and non-responders based on the standard error of measurement. While a subset of individuals exhibited consistent inhibitory responses, others displayed facilitatory or mixed patterns across sessions which is in line with previous studies [5; 22]. This also aligns with prior research suggesting that CPM responses can be influenced by state-dependent factors such as stress, mood, attention, and environmental context [25; 34]. Similarly, TSP measures were subject to substantial intersession variability, likely reflecting transient changes in central nociceptive processing between sessions [7].

To further explore variability across paradigms in the same participants, we introduced the NSCI measure. By normalising changes between sessions relative to the variability observed in the first session, NSCI provides a standardised metric for assessing consistency (i.e. participants with values close to zero showing highest consistency across sessions). Our analysis revealed that OA measures were the most stable, with NSCI values closest to zero, while CPM and TSP exhibited greater volatility. This suggests that OA, which despite having cortical influences over its brainstem circuitry [29; 42], appears to show more robust and stable responses across sessions compared to CPM. The anatomical and pharmacological differences between CPM and OA have been well documented [2], and our psychophysical data provide further evidence that they are governed by distinct top-down pathways that show different inherent variability. Further to this, we have demonstrated how NSCI can be used as a tool for making comparisons across different psychophysical paradigms within the same individuals. Using this approach, it may also be possible to profile changes across distinct endogenous pain modulation systems in chronic pain patients over time, which could help to identify specific dysfunctional central pathways to be targeted therapeutically [2].

We also introduced a CPM response status measure, derived from the integration of SEM thresholds, which we used alongside NSCI to help distinguish brain-based mechanisms underlying state- and trait-like CPM responses, an important feature of endogenous pain modulation systems [24]. For instance, individuals with consistent strong inhibitory or facilitatory responses (close to zero NSCI and CPM status close to +/-4) may exhibit robust and stable connectivity patterns within the descending pain modulation network. Participants with variable responses (high or low NSCI and close to zero CPM status) likely reflect state-like modulation, which have been suggested to be driven by fluctuations in sleep, stress, personality traits, expectation or mood [3; 8; 11; 36; 41].

By applying spectral DCM to rs-fMRI data, we then identified distinct patterns of effective connectivity associated with CPM response status. This aligns with prior research emphasising the importance of integrating neuroimaging with psychophysical measures to understand pain modulation mechanisms and variability [35]. Others have shown that efficient CPM responses measured in a single test session were associated with increased resting state functional connectivity between the PAG and key pain processing regions (e.g. primary and secondary somatosensory, motor, premotor, and dorsolateral prefrontal cortices) [15]. It has been also reported that functional connectivity within the descending pain modulation network was only present when participants demonstrated an inhibitory CPM profile and that faciliatory CPM was more evident in females [27]. Similar regions have also been demonstrated to be active during evoked response fMRI and functional near infrared spectroscopy imaging studies, which also showed activation of the ACC and insula cortices during CPM paradigms [33; 40]. Further, PAG connectivity is associated with variability within a single session of recorded CPM responses [14]. Here, we expand on these results and we illustrate using spectral DCM applied to rs-fMRI data that variability in CPM status across repeated test sessions was characterised by bottom-up excitatory influences from the PAG to the left AI, bilateral dACC, and right thalamus, as well as top-down inhibitory influences from the right AI to the thalamus and PAG. Taken together, these data suggest that stable or dynamic CPM responses over repeated test sessions may arise from the strength of effective connectivity between the PAG and other key cortical regions implicated in the descending modulation network.

We observed a significant positive correlation between PAG-to-right-AI effective connectivity and CPM facilitation. Specifically, participants with excitatory effective connectivity in this pathway exhibited consistent CPM facilitation, while those with inhibitory effective connectivity showed consistent CPM inhibition across sessions. This finding underscores the established role of the PAG as a central node in descending pain modulation [23; 28; 43], highlighting how bidirectional interactions between these regions can influence endogenous pain modulation. The PAG integrates nociceptive input both to and from the spinal cord and conveys modulatory signals to cortical regions such as the AI, which is involved in cognitive-affective appraisal and salience processing of pain [44; 46; 47]. The AI has been implicated in integrating interoceptive signals with emotional and attentional components of pain, making it a key region for top-down and bottom-up modulation of pain signals [4; 24; 38]. Increased excitatory connectivity between the PAG and AI may amplify the salience of pain signals, potentially explaining why some participants exhibit consistent heightened CPM responses in psychophysical paradigms. Conversely, inhibitory connectivity in this pathway might reflect enhanced top-down control, enabling consistent inhibition of nociceptive signals during CPM. These findings emphasise the importance of investigating effective connectivity within key pain-related networks to understand individual differences in pain modulation across repeated test sessions.

Whilst these measures give us insight into whether participants demonstrate stable (trait) or dynamic (state) like behaviour in endogenous pain modulation systems, longitudinal studies investigating the stability of these responses over extended periods could provide further insights into their trait-versus state-like characteristics. Establishing statistically sound response status criteria for TSP and OA, like the ± 2SEM criteria used for CPM, would enable us to investigate brain patterns underlying variability on these measures and elucidate underpinning neural mechanisms using the same approach as what we did with CPM. This is, however, outside the scope of this manuscript.

A limitation of this study is the relatively small sample size used in the DCM analysis. While DCM can provide robust insights into effective connectivity within the descending pain modulation network, smaller sample sizes can limit the generalisability of findings and the statistical power to detect subtle connectivity patterns. The use of advanced Bayesian frameworks, such as parametric empirical bayes, mitigates this issue to some extent by optimising parameter estimation and accounting for individual variability [9; 51]. However, larger sample sizes are essential to validate these findings and to explore potential subgroup differences in connectivity profiles during chronic pain.

In summary, this study provides new insights into the variability of endogenous pain modulation across repeated test sessions and sheds light into how this relates to effective connectivity in descending pain modulation systems. The introduction of a novel NSCI and CPM response status measure also offers a new framework for quantifying consistency within descending pain controls systems in the same participants. By integrating psychophysical measures with resting state functional MRI, we also demonstrated the potential of spectral DCM analysis to elucidate the mechanisms driving intersession variability in CPM responses. These findings highlight the importance of combining behavioural and neuroimaging data to advance our understanding and profiling of endogenous pain modulation dynamics over repeated test sessions.

## Acknowledgements

This study was funded through an Academy of Medical Sciences Springboard grant (SBF007\100108). The analysis plan for this study (excluding exploratory analyses) was preregistered and available in Open Science Framework (OSF) Registries on the following link https://osf.io/x9uf8. Preregistration includes study design, variables, and treatment conditions and description of the analysis plan (including specification of the sequence of analyses reported).

